# Soil inoculum identity and rate jointly steer microbiomes and plant communities in the field

**DOI:** 10.1101/2021.08.22.456585

**Authors:** Xu Han, Yingbin Li, Yuhui Li, Xiaofang Du, Bing Li, Qi Li, T. Martijn Bezemer

## Abstract

The importance of soil inoculation to engineer soil microbiomes and ultimately entire ecosystems is becoming widely acknowledged. Inoculation with soil from different ecosystems can induce directional changes in soil and plant communities and promote the restoration of degraded ecosystems. However, it is unknown how such inoculations influence the soil microbiome, how much inoculum is needed, and whether inocula collected from similar ecosystems will steer the microbiome in different directions. We conducted a three-year soil inoculation field experiment at a degraded grassland and used two different soil inocula both from grasslands with three inoculation rates. Our results show that inoculation with soil that originates from different donor grasslands steers the soil microbiome as well as the plant communities at the inoculated site which was a degraded grassland into different directions and that these effects were stronger with increasing amount of soil used to inoculate. Inoculation with upland meadow soil introduced more keystone genera and resulted in more complex biotic networks in the soil than inoculation with meadow steppe soil. Our experiment highlights that soil inoculation can steer soil microbiomes in the field and that the direction and speed of development depend on the origin and the amount of soil inoculum used.

## Introduction

Grasslands cover about 40% of the earth’s surface, providing a large number of ecosystem services [1, 2]. However, the intensification of land use has resulted in a worldwide degradation of species-rich forage grasslands and undermined their ability to support biodiversity and ecosystem functioning [3–5]. Recent studies have shown that inoculation with soil from a late successional stage can alter soil microbial communities and facilitate the establishment and growth of late-successional plant species [6–9]. This is particularly efficient when the inoculation is carried out after the top-soil layer has been removed [8, 10]. For example, Wubs *et al.* recently showed that in a former arable field the composition of both the soil and the plant community was steered towards a predefined target community when using soil inoculum collected from a semi-restored grassland and from a heathland [8]. While there is now evidence that inoculation with soil from two distinctly different ecosystems can induce directional changes in the soil and in plants communities in the field [11], a major outstanding question is how specific these inoculations are and whether inoculation with soil originating from underneath different plant communities but from a similar ecosystem will also lead to differential development of the soil microbial community and ultimately the entire ecosystem at the recipient site.

In empirical soil inoculation studies, so far, greatly varying amounts of donor soil have been used ranging from 0.016 cm up to 40 cm layers of donor soil [12, 13]. We may expect that there is a positive relationship between the establishment of soil biota and plants and hence the success of the inoculation, and the amount of soil that is used. Results from a controlled pot experiment show that indeed there was a positive effect between the amount of inoculated soil and plant biomass [14]. However, in the field, and over longer time periods, it is also possible that due to positive feedbacks between the soil and plant communities, inoculation with a small amount of donor soil will give similar “steering power” to the soil microbiome and plant communities as larger amounts of donor soil [8, 11]. How the amount of donor soil that is used for inoculation influences the establishment of soil microbiome and plant communities in the field is still not known. In restoration projects, the donor soil typically originates from a nature area, and collecting this soil will inevitably cause some destruction of the donor area. Hence it is essential to better understand the minimum amount of soil inoculum required for successful restoration to avoid over-exploitation of the donor area.

We conducted a soil inoculation experiment in a degraded grassland and measured the development of the soil microbiome, of soil nematodes and of the plant communities over a period of three years following soil inoculation. In soil from the experimental fields we determined the bacterial and fungal communities and identified the nematode communities. We used two donor grasslands: a meadow steppe and an upland meadow, and inoculated three different amounts of donor soil in experimental plots that were established after removing the top soil layer at the degraded site (Fig. 1). We hypothesized that inoculation with the different donor grassland soils would result in different soil microbiomes and plant communities; and that the effect of soil inoculation would increase with increasing amounts of donor soil used to inoculate the field plots.

**Figure 1.**
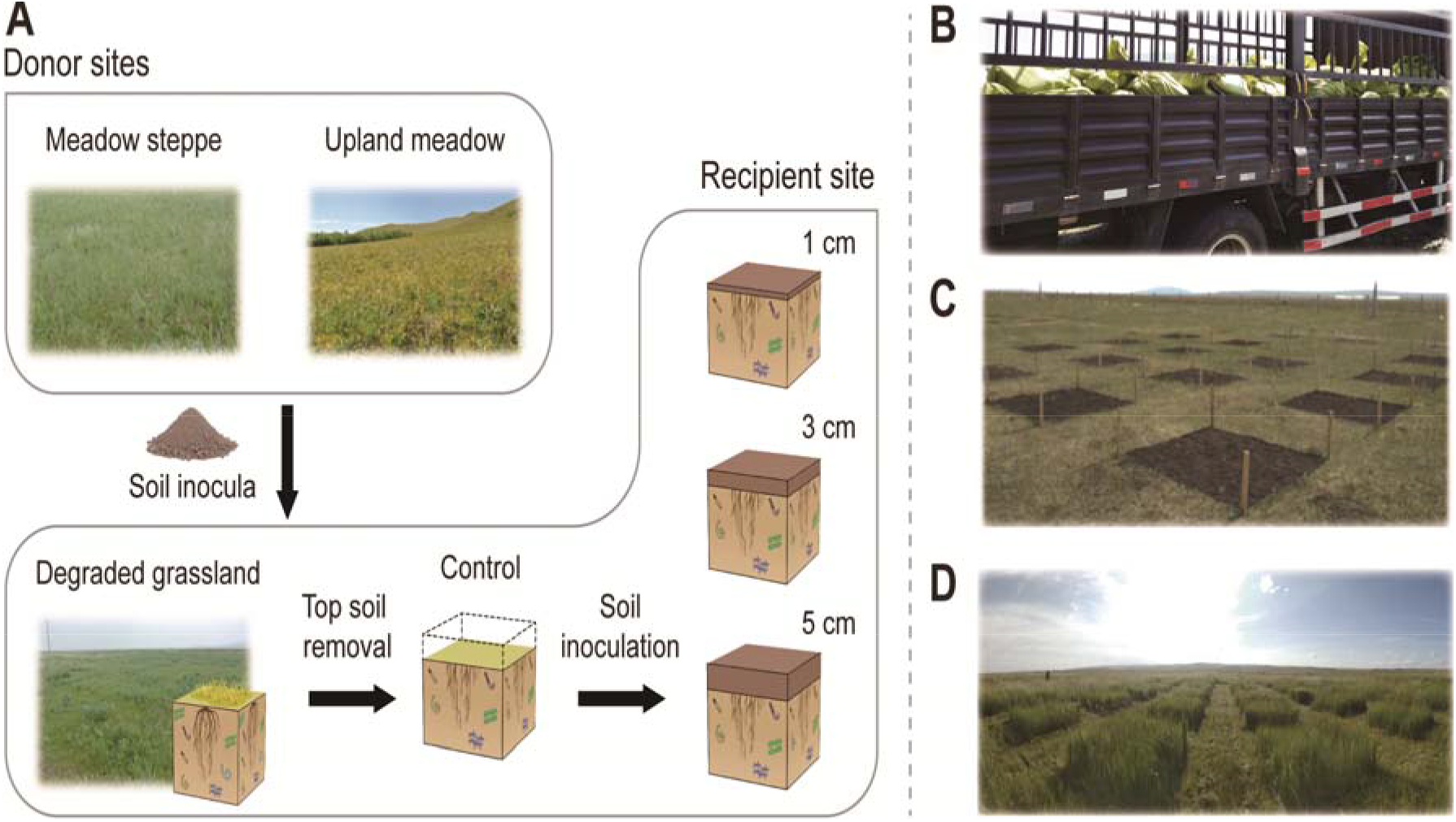
Graphical illustration of the experimental design. **A**, Three replicate meadow steppes and upland meadows were selected as donor sites. At each donor location, the soil of the top 10 cm was excavated, homogenized and transported to the recipient site. A degraded grassland was selected as recipient site. The top 5 cm of the soil was removed before inoculation. The experiment was set up with three fully replicated blocks. In each block, one plot without inoculation (TSR) was also established. We established 19 experimental plots of 2 × 2 m^2^ (2 soil types × 3 replicate sites × 3 inoculum amounts + TSR) in each block and a total of 57 plots. **B**, the transfer of soil inoculum in 2018 to the field; **C**, the experimental field site after inoculation in 2018; **D**, and three years later, in 2021.

## Methods

### Experimental design

The experiment was carried out at a degraded grassland near the Erguna Forest-Steppe Ecotone Research Station of the Chinese Academy of Sciences (50° 10′ 46.1″ N, 119° 22′ 56.4″ E). The mean annual temperature is −2.4 °C with minimum value in January (−28 °C) and maximum value in July (19.1 °C). The mean annual precipitation is 361.6 mm mainly concentrated in summer and autumn. The soil type is classified as chernozem in the World Reference Base for Soil Resources (WRB) [15].

In May 2018, a degraded grassland was selected as recipient site. Three replicates of meadow steppe and upland meadow were selected as donor sites. The top 5 cm of the soil from the degraded grassland was removed (top soil removal, TSR). In the experiment, we inoculated soil from two donor grassland ecosystems, a nearby meadow steppe and an upland meadow. The meadow steppe was a grassland with moderate disturbance of human activity due to mowing each year dominated by xerophytic grasses. The upland meadow belongs to the forest-steppe ecotone with little disturbance dominated by mesophytic forbs. Three replicate sites were set for each donor site with about a 1 km distance between each. At each donor location, the top 10 cm of soil was excavated and homogenized and inoculated in plots within a week. At the recipient site the soil was inoculated at three inoculation amounts (1 cm, 3 cm and 5 cm) with a rate of 1.34 g cm^−3^ for meadow steppe and 1.13 g cm^−3^ for upland meadow, respectively (based on corresponding bulk density). The experiment was set up with three replicated blocks. In each block, one plot with top soil removal but without inoculation (TSR) was also established. We established 19 experimental plots (2 soil types × 3 replicate sites × 3 inoculum amounts + TSR) in each block and a total of 57 plots. Each plot was 2 × 2 m^2^ in size with 2 m wide paths between them. The central 1 × 1 m^2^ quadrat was used for vegetation recording and small quadrats (0.25 × 0.25 m^2^) were set adjacent to the central quadrat at a different locations each year for destructive sampling of soil and plant properties. In 2020 we also collected soil samples at the donor sites and recorded the biomass and vegetation composition.

### Soil sampling and analysis

In August of 2018, 2019 and 2020, within each plot, ten soil cores (2.5 cm diameter, 10 cm deep) from one of the adjacent quadrats (0.25 × 0.25 m^2^) were collected and pooled as a composite sample for each plot. After gentle homogenization and removal of roots, 10 g fresh weight of soil was immediately stored at −80 °C for DNA analysis. About 100 g fresh weight of soil was kept in a plastic bag at 4 °C for nematode community analysis, and the remaining soil was sieved through a 2 mm mesh, air dried and analyzed for soil physicochemical properties. Soil moisture was determined by oven-drying subsamples at 105 °C for 24 h. A soil: water (1:2.5) mixture was used for measuring soil pH with a glass electrode. Total carbon (TC) and total nitrogen (TN) content in each sample were determined using a TruSpec CN Elemental Analyzer (Leco Corporation, USA). Total phosphorus (TP) was determined by the method of molybdenum-antimony colorimetric using a spectrophotometer (Shimadzu Inc., Kyoto).

### DNA extraction and amplicon sequencing

We extracted bacterial and fungal DNA from 0.5 g soil using the FastDNA™ Spin Kit for Soil (MP Biomedicals, USA), followed the manufacturer’s protocol. Bacteria was amplified using the primers 515F (5′- GTGCCAGCMGCCGCGG -3′) and 907R (5′- CCGTCAATTCMTTTRAGTTT -3′), and fungi was amplified using the primers ITS86F (5′- GTGAATCATCGAATCTTTGAA -3′) and ITS4R (5′- TCCTCCGCTTATTGATATGC -3′). PCR reactions were conducted with 10 ng of extracted soil DNA as template per 20 μl PCR reaction which contained 4 μl 5 × FastPfu buffer, 2 μl 2.5 mM dNTPs, 0.4 μl FastPfu Polymerase, 0.2 μl BSA, 1.8 μl ddH20 and 0.8 μl of each forward and reverse primers. PCR cycling conditions were set for 95 °C for 3 min, followed by 27 cycles for bacteria and 35 cycles for fungi of denaturation at 95 °C for 30 s, annealing at 55 °C for 30 s, extension at 72 °C for 45 s and a final extension of 72 °C for 10 min. Amplicons were sequenced on the Illumina MiSeq PE300 platform. Sequences were quality trimmed using FASTP v0.19.6, merged with FLASH v1.2.11, dereplicated and clustered to OTUs with UPARSE v7.0.1090 according to 97 % similarity. For bacteria, we assigned the sequences to taxonomic groups using the SILVA database and we assigned fungi sequences to taxonomic groups using the UNITE database. For all following calculations, OTUs that occur less than 3 times with relative abundances < 0.01% were disregarded. To account for differences in sequencing depth, we rarefied the bacteria and fungi to 19,067 and 59,331 reads, respectively. For sequencing, we merged the soil of three blocks for each treatment in the first year. So, a total of 21 samples (2 soil types × 3 replicate sites × 3 inoculum amounts + 3 TSR) were analyzed in 2018. In 2019 and 2020, all 57 samples were sequenced.

### Nematode community analysis

Soil nematodes were extracted from 100 g of fresh soil according to a modified cotton-wool filter method [16, 17] and fixed in 4 % formaldehyde solution. We counted the total number of nematodes and at least 100 individuals were identified to genus level [18–20].

### Plant community analysis

In August of 2018, 2019 and 2020, species richness (number of species) and cover of each plant species were recorded in the 1 × 1 m^2^ central quadrat of each plot. The cover was represented by estimating the percentage of projected area of all plant species within the central quadrat. The aboveground vegetation was harvested by clipping all biomass at 1 cm from the soil level in one of the adjacent quadrats in each plot. In the third year belowground biomass was also recorded. Three soil cores (5 cm diameter, 10 cm deep) inside the adjacent quadrat were collected from each plot and weighed, and roots were cleaned with water and a 2 mm sieve. All aboveground and belowground biomass were oven-dried at 60 °C to constant weight and dry biomass was recorded.

Plant species were classified as annual or perennial according to the Flora of China [21]. The ratio of perennial cover to total cover was calculated to show the effect of grassland restoration [22]. From the plant species database, we further classified indicator species for degenerated sites, meadow steppe and upland meadow based on the Flora of China [21]. *Leymus chinensis* was considered as the most important restoration target species at the middle stages of restoration [23].

### Data analysis

All statistical analyses were carried out in R version 3.6.1. Composition of the vegetation was analyzed based on the relative plant cover. We analyzed univariate response data using linear mixed models, with separate models for soil physicochemical properties, soil biota and plant community. First, we assessed the overall effect of year, soil type, inoculum amount and their interactions by linear mixed models with repeated measures, with site replicate and block as random factors. Second, we assessed the differences between all inocula and control treatment each year using Post hoc Scheffe tests with the glht function in the multcomp package. Three linear mixed models, for each year, were constructed respectively with block as random factors. Linear mixed models were constructed by lmer (in the lme4 package) function respectively [24].

For 2020, we tested the effect of soil inoculation by calculating the RR (response ratio) between treatments and control, donor meadow steppe or donor upland meadow by

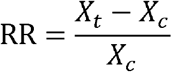

Where Xt is the value of variables in soil inoculation treatments. Xc is the value of corresponded variables of control, donor meadow steppe or donor upland meadow in the same block or same site replicates. Linear mixed models were used to test the soil type and amount effect on RR with soil type, amount and interaction as fixed effects, and block and site replicates as random factors.

To visualize the effect of year, soil type, inoculum amount and their interactions on soil microbiomes, nematodes and plant communities, we performed unconstrained principle coordinates analysis (PCoA) and permutational analysis of variance (PERMANOVA). Plant cover was Hellinger-transformed before this analysis. Separate permutational analysis with different years were conducted to test the year effect on soil organisms and plant communities. To examine to what extent the composition of the soil microbial and nematode communities in the inoculated plots resembled those of the donor site, we conducted PCoA with the third-year data from the experimental plots together with the donor sites. Ordination analyses were performed using the R package phyloseq based on Bray-Curtis dissimilarities and permutational analysis of variance using the functions adonis2 in the vegan package [25, 26].

We subsequently compared the similarity between inoculated plots and control and donor sites. We calculated the pair-wise similarity between each treatment to control, donor meadow steppe and donor upland meadow by 1– Bray-Curtis dissimilarity. We also assessed soil inoculation effects on similarity using a linear mixed models, soil type, inoculum amount and interaction as fixed effects, block and site replicate as random factor.

We single out the significantly increased genera (relative abundance) in meadow steppe and upland meadow inoculation treatment than control plots which were considered to be directly introduced by soil inoculation or indirectly promoted by inoculation. Those genera confirmed by both indicator species (in the indicspecies package) and LRT (likelihood ratio tests, in the edgeR package) analysis [27].

We constructed co-occurrence networks by spearman rank correlations between all pairs of bacteria, fungi and nematode genera, significant correlations (r > 0.6 and *P* < 0.001) were shown. All networks were visualized with the Fruchterman-Reingold layout with 10^4^ permutations in igraph. The network complexity was calculated as linkage density. Greater complexity was associated with multiple functions of microbial communities [28]. We generated random networks compare to our networks to test our networks were not random by erdos.renyi.game function in igraph [29]. Microbial taxa that frequently co-occur with other taxa in microbial co-occurrence networks are thought to be ecologically important, so the 10 keystone genera with the highest node degrees of co-occurrence were selected [27, 30–32]

## Results

### Inoculum identity and amount steer soil microbiomes and nematode communities

The identity of the donor soil and inoculation amount both significantly affected the soil microbiomes and nematode communities in the recipient plots (Fig. 2; Table. 1). Already after three years, the bacterial and fungal communities had diverged towards those of the donor soil and inoculum amount strengthened this divergence (Fig. 2). Soil inoculation effects on the nematode community were limited in the first year but significant in the second and third year (Table. S1). Richness of bacterial and fungal communities were also affected by soil inoculation (Fig. S1, Fig. S2). The amount of inoculum had a positive effect on fungal richness (F = 15.26, *P* < 0.001) but a negative effect on bacterial richness (F = 3.21, *P* = 0.044). Additionally, richness of the bacterial community was lower in soil inoculation with upland meadow than that with meadow steppe (F = 5.6, *P* = 0.02). Together, these results show that donor soils originating from a similar ecosystem can steer the soil communities at the recipient site towards the specific donor community but also, for the first time that in the field, this is accelerated when increasing amounts of inocula are used.

**Figure 2.**
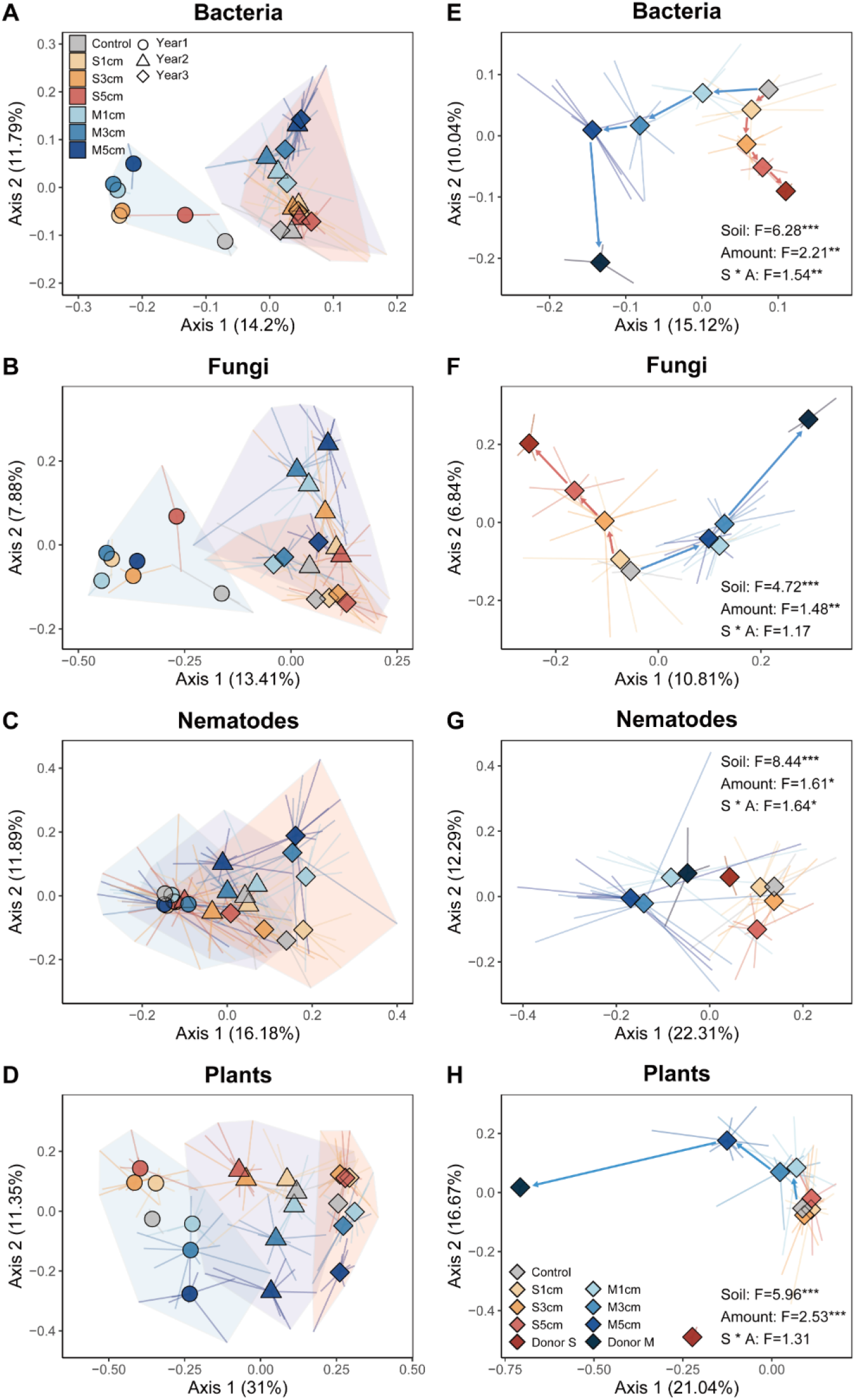
Effects of soil inoculation on soil bacteria, fungi and nematodes, and plant communities. Separate PCoA ordinations using Bray-Curtis distance are presented in three years (**A**, **B**, **C** and **D**) and the third years (**E**, **F**, **G** and **H**). Percentage of variation given on each axis refers to the explained fraction of total variation. The colors depict soil inoculation treatments: Control, the top soil removed treatment (TSR); S 1cm, meadow steppe soil inoculation with 1 cm depth; S 3cm, meadow steppe soil inoculation with 3 cm depth; S 5cm, meadow steppe soil inoculation with 5 cm depth; M 1cm, upland meadow soil inoculation with 1 cm depth; M 3cm, upland meadow soil inoculation with 3 cm depth; M 5cm, upland meadow soil inoculation with 5 cm depth; Donor S, donor meadow steppe; Donor M, donor upland meadow. The different shapes and shaded areas jointly represent plots in different years: circle, for the first year; triangle, for the second year and rhombus, for the third year. The result of PERMANOVA for three years were showed in table 1 and for the third year were listed above, * *P* < 0.05; ** *P* < 0.01; *** *P* < 0.001.

**Table 1.** Adonis test of the effect of year, soil type, inoculum amount and their interactions on soil and plant communities.

|  | Bacteria |  | Fungi |  | Nematodes |  | Plants |  |
| --- | --- | --- | --- | --- | --- | --- | --- | --- |
|  | F | *P* | F | *P* | F | *P* | F | *P* |
| Year | 13.4 | <0.001 | 11.89 | <0.001 | 18.24 | <0.001 | 71.64 | <0.001 |
| Soil | 10.53 | <0.001 | 7.33 | <0.001 | 5.41 | <0.001 | 19.48 | <0.001 |
| Amount | 3.55 | <0.001 | 2.09 | <0.001 | 1.91 | 0.008 | 6.7 | <0.001 |
| Year * Soil | 1.1 | 0.305 | 1.13 | 0.237 | 4.07 | <0.001 | 4.67 | <0.001 |
| Year * Amount | 0.89 | 0.656 | 0.63 | 0.998 | 1.21 | 0.11 | 2.17 | 0.011 |
| Soil * Amount | 2.07 | <0.001 | 1.51 | 0.016 | 1.35 | 0.131 | 4.18 | <0.001 |
| Year * Soil * Amount | 0.84 | 0.786 | 0.69 | 0.992 | 0.97 | 0.417 | 1.01 | 0.432 |

### Soil inoculum identity and amount steer plant communities

Soil inoculation also significantly affected plant community composition (Fig. 2; Table. 1) and steered the composition of plant communities towards the specific grassland donor sites (Fig. 2). The proportional cover of perennials increased strongly over time (F = 168.82, *P* = 0.006) and was higher in plots inoculated with upland meadow soil than in plots that received meadow steppe soil (F = 92.57, *P* < 0.001) (Fig. S3). With increasing amount of inoculum, species richness of the plant community also increased and plant species richness was higher in upland meadow than in meadow steppe inoculated plots (Fig. S2). When compared to the uninoculated control (top soil removed only, TSR), soil inoculation suppressed the growth of degenerate indicator species, and increased the relative cover and abundance of the target species *L*. *chinensis* (Fig. S3). The total cover of all target species increased over time (meadow steppe: F = 92.4, *P* = 0.011; upland meadow: F = 79.37, *P* = 0.012) but this was not affected by inoculum amount (Fig. S4). More upland meadow target species colonized in plots inoculated with upland meadow soil (F = 15.62, *P* < 0.001) but meadow steppe target species did not colonize more successfully in plots inoculated with meadow steppe soil than in control plots (Fig. S4). In the third year, belowground (root) biomass was significantly higher in inoculated than in control plots, and this effect increased with increasing amount of inoculum used but no significant effect was observed for aboveground biomass. Concentrations of soil nutrients and soil moisture also increased with increasing inoculum amount (Fig. 3, Fig. S5).

**Figure 3.**
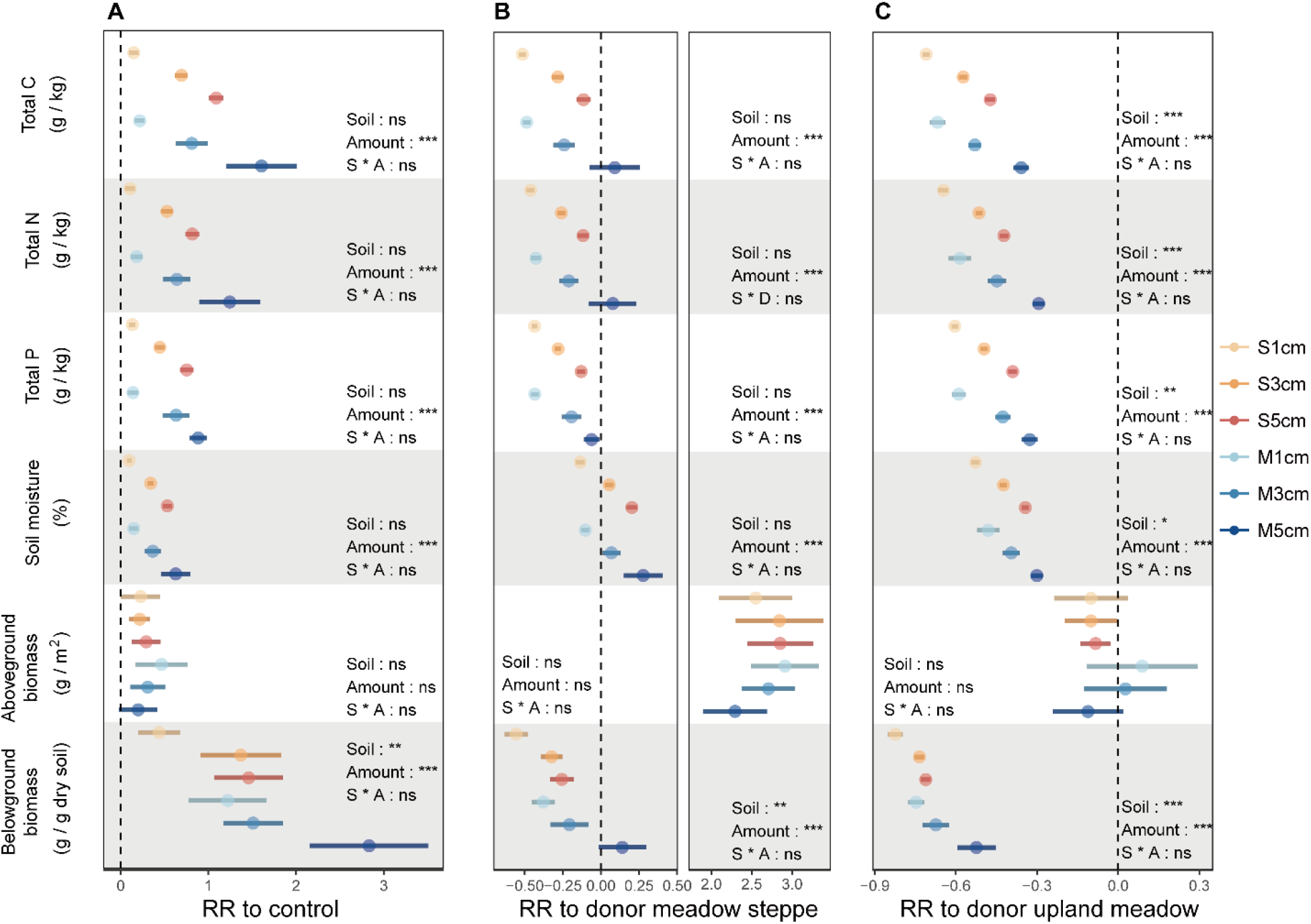
Response ratio of soil and plant traits to control, donor meadow steppe and donor upland meadow. Error bars represent ± SE. The category definitions are as follows: S 1cm, meadow steppe soil inoculation with 1 cm depth; S 3cm, meadow steppe soil inoculation with 3 cm depth; S 5cm, meadow steppe soil inoculation with 5 cm depth; M 1cm, upland meadow soil inoculation with 1 cm depth; M 3cm, upland meadow soil inoculation with 3 cm depth; M 5cm, upland meadow soil inoculation with 5 cm depth. * *P* < 0.05; ** *P* < 0.01; *** *P* < 0.001.

### Similarity to the control and the donor sites

We further analyzed the effectiveness of inoculation by comparing the similarity of the soil and plant communities of inoculated and uninoculated plots and the similarity of inoculated plots to the two donor sites in the third year (Fig. 4). With increasing amount of soil inoculum, inoculated plots became more dissimilar from the control (uninoculated) plots, and this was particularly so for bacterial communities and for plants. Inoculation with meadow steppe soil increased the similarity to this donor more than inoculation with upland meadow for bacterial, fungal and nematode communities and a positive effect of inoculation amount was found for bacterial and fungal communities. Similarly, inoculation with upland meadow soil resulted in higher similarity to the upland meadow donor site for fungi and plants and a positive effect of the amount of inoculum was found for bacterial, fungal and plant communities.

**Figure 4.**
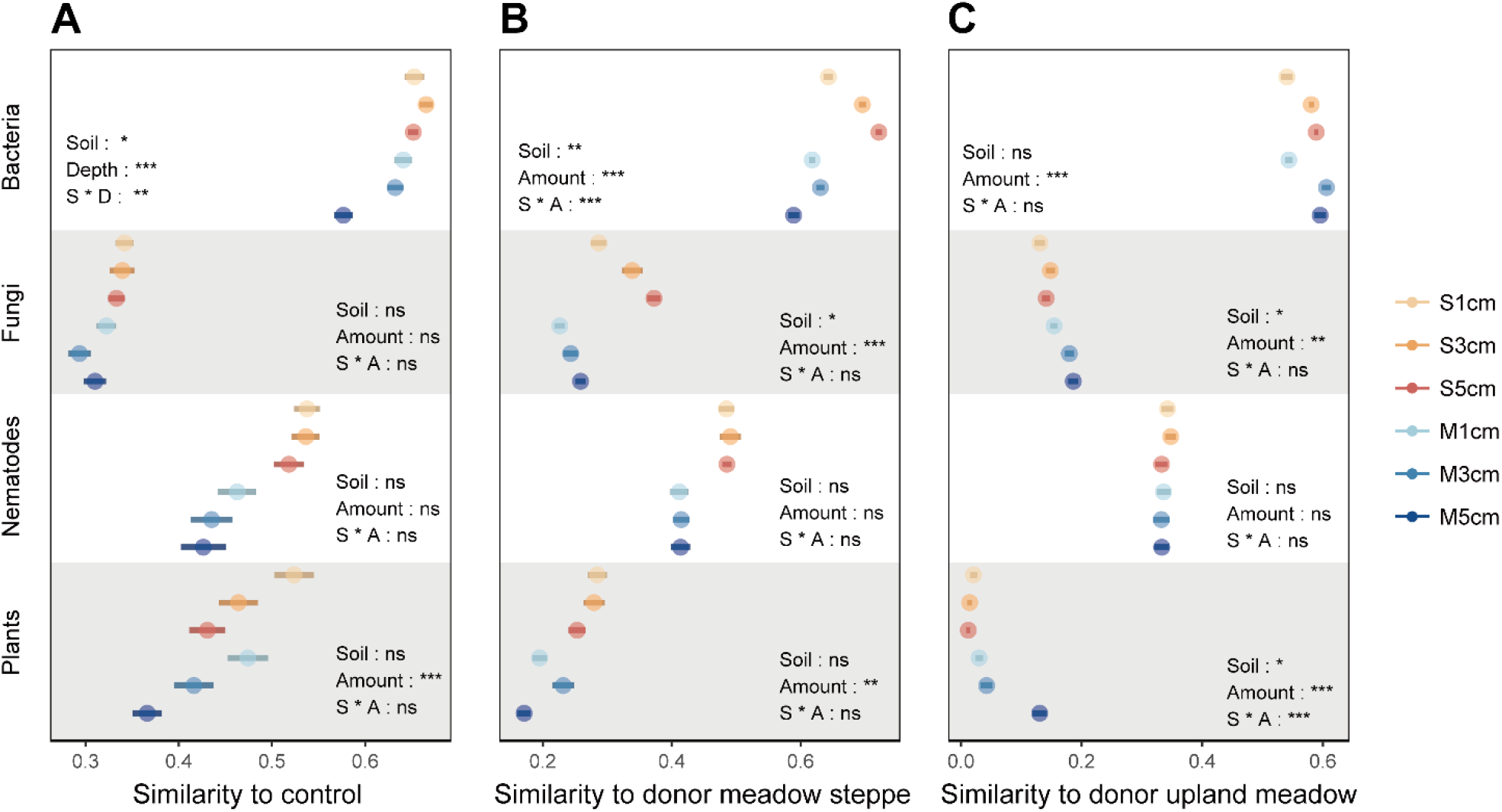
Similarity between soil inoculation treatments to TSR or donor treatments. Error bars represent ± SE. The category definitions are as follows: S 1cm, meadow steppe soil inoculation with 1 cm depth; S 3cm, meadow steppe soil inoculation with 3 cm depth; S 5cm, meadow steppe soil inoculation with 5 cm depth; M 1cm, upland meadow soil inoculation with 1 cm depth; M 3cm, upland meadow soil inoculation with 3 cm depth; M 5cm, upland meadow soil inoculation with 5 cm depth. * *P* < 0.05; ** *P* < 0.01; *** *P* < 0.001.

### Soil networks

Soil genera with higher relative abundance in the inoculation plots than in the control were selected (Fig. 5). These genera may be directly introduced by soil inoculation or indirectly promoted following inoculation. To confirm whether these genera play an important role in the soil community, we explored the distributions of bacterial, fungal and nematode communities in co-occurrence networks (Fig. 6A, B). The ten keystone genera at the center of the network (higher node degree values, i.e. related with more other genera) were selected (Fig. 6C). Most of those genera significantly correlated with soil and plant traits (Fig. 6D). We also found that eight out of these ten keystone genera increased after inoculation with upland meadow soil, and they also positively correlated with the amount of soil inoculum. However, the genera that increased after inoculation with meadow steppe soil did not occupy the central position in the biotic network. Network complexity was also higher after inoculation with upland meadow (1.86) than after inoculation with meadow steppe soil (1.55) (Table. S2).

**Figure 5.**
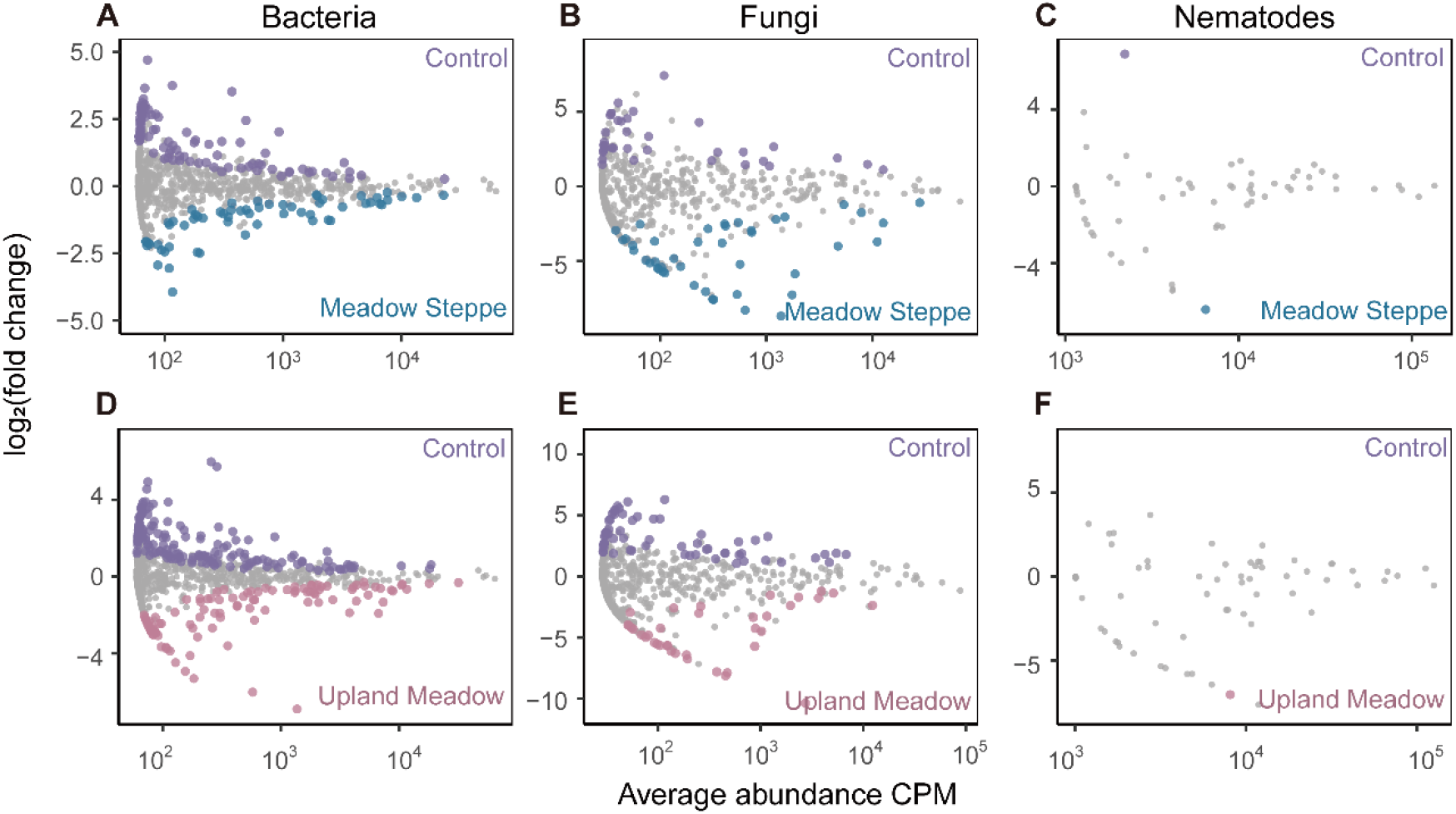
Soil inoculation effects on specific sets of bacteria, fungi and nematodes. Abundance patterns of meadow steppe soil inoculation treatments in bacterial (**A**), fungal (**B**) and nematode (**C**) communities (blue) and of upland meadow soil inoculation treatments in bacterial (**D**), fungal (**E**) and nematode (**F**) communities (pink) in comparison with the control (purple). The X-axis shows log_2_ (fold change), and the Y-axis shows –log_10_ (P value) (control relative to meadow steppe inoculation or upland meadow soil inoculation). Blue, pink and purple points showed specific genera in meadow steppe, upland meadow and control respectively, and non-differential genera are colored in gray (likelihood ratio test, *P* < 0.05, FDR corrected).

**Figure 6.**
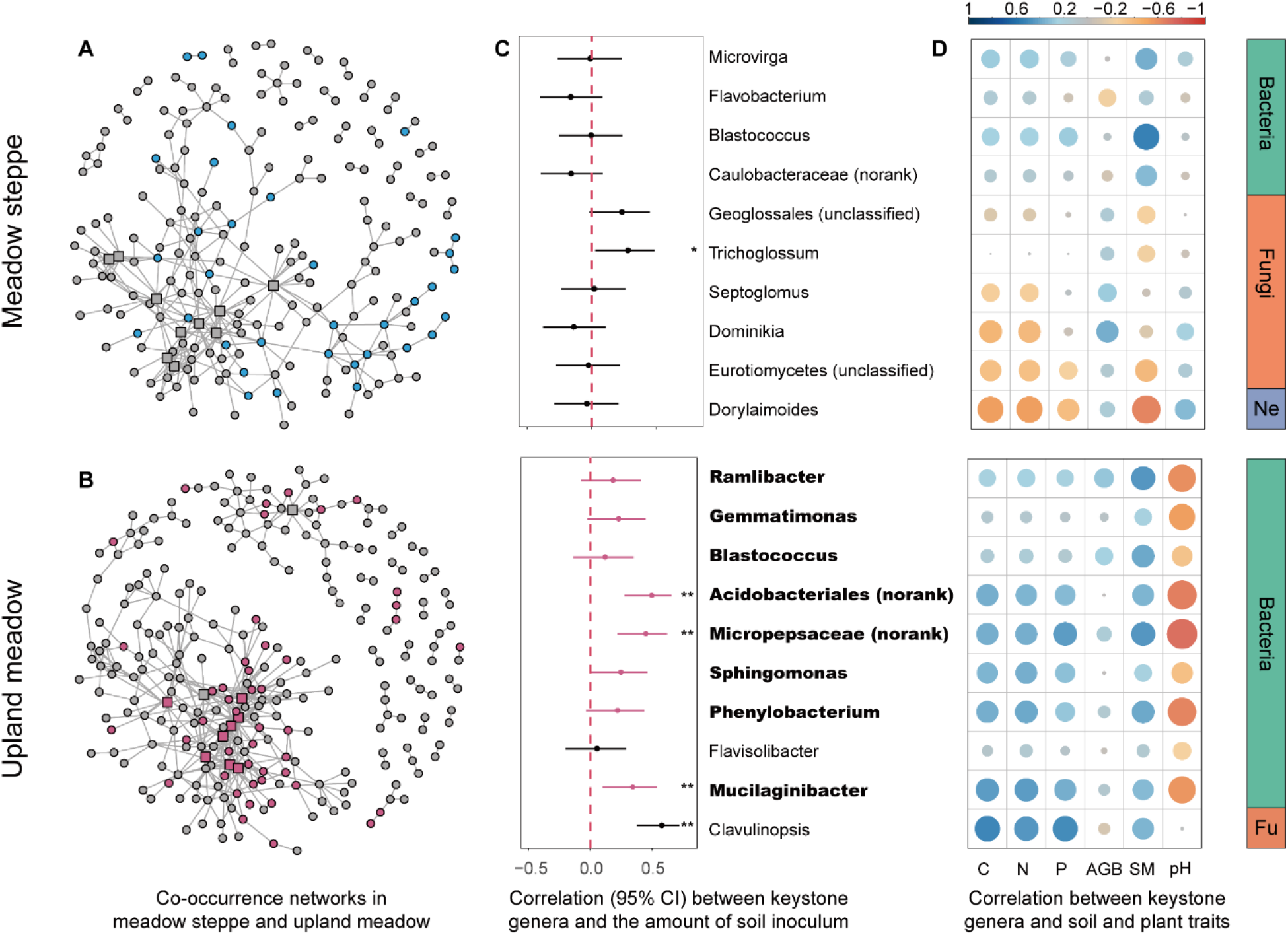
The network analysis and the increased genera after inoculation with different donor soil. Co-occurrence networks at genus level based on meadow steppe (**A**) and upland meadow (**B**) soil inoculation. Correlation between the relative abundance of 10 keystone genera with the highest node degree and soil inoculum amount (**C**) and soil and plant traits (**D**). Blue (in **A**) and pink points (in **B**) indicated the genera significantly increased after inoculation meadow steppes and upland meadow soil than control plot. Square in networks (in **A** and **B**) and bold genera names (in **C**) indicated the genera significantly increased after soil inoculation. * *P* < 0.05; ** *P* < 0.01; *** *P* < 0.001. C: soil total carbon, N: soil total nitrogen, P: soil total phosphorus, AGB: aboveground biomass, SM: soil moisture, Ne: nematodes, Fu, fungi.

## Discussion

This soil inoculation experiment at a degraded grassland with two different soil inocula and three inoculation rates shows that in the experimental plots soil and plant communities diverged and that these effects were stronger when more soil was used for inoculation. Moreover, inoculation with upland meadow soil introduced more keystone genera that may play important role in the soil networks than inoculation with meadow steppe soil. These findings emphasize that there are specific effects of donor soil on the developing soil microbiomes and on the wider above- and belowground communities and that these effects depend on the identity of the donor even when comparing two grassland donors.

The importance of the role of soil organisms in the restoration of plant communities is becoming widely acknowledged now [33, 34]. Soil inoculation can be used to promote the colonization of soil organisms at the start of restoration. This study now shows that inoculation with soil communities that originate from different donor grasslands steers soil microbiomes as well as plant communities of the inoculated degraded grassland site into different directions. This result highlights the importance of selecting the correct donor site in restoration projects. In our study, inoculation with both meadow steppe and upland meadow soil led to a higher similarity to the specific donor but plant community inoculation with upland meadow soil caused much stronger effects than inoculation with meadow steppe soil. More target plant species from the upland meadow colonized the inoculated site after inoculation with upland meadow soil, while colonization of meadow steppe target plant species after inoculation with meadow steppe soil was less successful. These findings suggest that there are distinct differences in colonization or establishment success of soil microorganisms and plant among different donor soils. Interestingly, these findings also show that differences in the initial composition of the whole soil inoculum, can alter interactions in the soil networks at the recipient site and this suggests that the choice of donor can have important consequences for the functioning of soil communities and ultimately for the functioning of the entire ecosystem that is restored. To what extend the inoculum-specific changes in the plant communities were directly caused by introducing different species of plants (e.g. as seeds or root pieces) in the inoculated soil or whether this was due to different rates of establishment of the plants mediated by the altered soil community requires further work.

An important result from this study is that the changes in the soil and plant communities and the similarity to the donor sites increased almost linearly with the amount of inoculum used. These findings support our second hypothesis that the effect of soil inoculation on soil microbiomes and plant communities is accelerated with increasing amounts of soil used for inoculation. In an experiment with trees, stronger inoculation effects were observed when higher amounts of soil inoculum were used on the height and diameter of ectomycorrhizal tree species [35]. It is likely that with the increasing amount of soil, the density of seeds, root pieces and soil organisms linearly increased and that this results in increased establishment of the introduced organisms [36]. However, it is important to note that for many plant species, germination of seeds from the seedbank only occurs at the soil surface [8] and in our experiment for all inoculum amounts the soil surface was covered with a layer of inoculated soil. Inoculum amount can also influence the establishment and survival of the introduced soil biota. It is likely that the introduced soil community survives and establishes best in the local habitat and that increasing the amount of inoculated soil at the recipient site increased resemblance to the original habitat conditions, e.g. by preventing the effects of desiccation and by better resembling the original abiotic conditions such as nutrient availability and pH [11, 12]. A better functioning new soil community could, in turn, be important for the establishment of target plant species. Many functions that soil biota provide are density dependent [14] and the density of soil organisms should be sufficiently high to avoid mortality and inbreeding depression [11, 12]. In this study, upland meadow soil inoculation also introduced some keystone taxa which may play key roles by determining microbiome functioning, such as, *Acidobacteriota*, *Proteobacteria* and *Bacteroidota*. The abundance of those genera positively correlated with the amount of inoculum and with the soil and plant traits. These bacteria are important in soil carbon and nitrogen cycling especially when nutrient availability is low [37–39], which may, in turn, influence the biogeochemical processes at the recipient site. Future experiments are needed to assess whether these keystone genera or inoculation sensitive genera directly or indirectly influence the other community members in the soil microbiome and the performance of plant communities.

Over the first three years following soil inoculation, the restoration of the degraded grassland soil and plant communities (i.e. the similarity to the donor sites) improved considerably. Importantly, the effects that we observed were not strongest immediately after inoculation and then declined, but instead the effects increased over time [35]. For example, soil inoculation effects on the nematode community were limited in the first year but significant in the second and third year. This clearly shows that the effects of inoculation were not remnants of what was left behind in the inoculated soil, but instead, that soil inoculation initiated the development of new soil and plant communities at the degraded site. Nematodes occur at different positions in the soil food web, and our results show that also higher trophic levels of the soil food web are influenced by soil inoculation, already two to three years after inoculation, even though organisms at these trophic levels are typically slow-growing and have low dispersal [40, 41]. These results show that whole soil inoculation is a promising method by which not only microbiomes but entire soil food webs can be molded rapidly in the field and as such can be a very useful tool in ecosystem restoration projects.

Via soil inoculation we created new soil and plant communities in the field. It is plausible that these new aboveground and belowground communities will interactively influence each other. A recent long term study in a species rich grassland on a former arable field showed that it can take up to a decade before such interactions become apparent [34]. Long term measurements are needed at the experimental field site where after three years both plant and soil communities changed considerably, to determine how the new aboveground and belowground communities will interact and how this will change the functioning of these grassland ecosystems and the composition of species in those ecosystems.

Overall, the experiment highlights that soil inoculation can promote restoration of degraded ecosystems but that the way and direction the ecosystem develops after inoculation depends strongly on the origin of the inoculum and the amount of inoculum used. Long-term measurements are needed to determine whether the ecosystems that received small amounts of inoculum, in the longer term, will exhibit a similar but slower pattern of development as the ecosystems that received larger amounts of inoculum or whether instead the former will become similar to the uninoculated controls over time. Such long term data will also reveal whether and for how long the ecosystems that were inoculated with different donor soils will continue to develop into different communities (e.g. via different aboveground-belowground interaction loops), or whether they will converge over time.

## Supporting information

Supplementary matrials

## Acknowledgments

We thank X.G. Han and Z.W. Wang for discussion and selection of the degraded grassland. We thank J. Guo for the help of vegetation investigation. Q.L. acknowledges funding from International cooperation program of Chinese Academy of Sciences (151221KYSB20200014) and National Natural Science Foundation of China (41877047). T.M.B. acknowledges funding from The Netherlands Organization for Scientific Research (NWO VICI grant 865.14.006).

## Author contributions

T.M.B and Q.L. designed the study. X.H. and Y.B.L set up the experiment and analyzed the data. X.H., Y.H.L, X.F.D., B.L. and Y.B.L collected the samples and conducted the measurements. X.H., Q.L. and T.M.B. wrote the first draft, and all authors contributed to the editing of the paper.

## Data and materials availability

All data needed to evaluate the conclusions in the paper are presented in the paper and/or the Supplementary Materials. Additional data related to this paper may be requested from the corresponding author.

## Competing interests

The authors declare that they have no competing interests.

## Notes

### Competing Interest Statement

The authors have declared no competing interest.

