## Supplementary matrials for "Soil inoculum identity and rate jointly steer microbiomes and plant communities in the field"

**Supporting Online Material:**

**Materials and Methods:**

***Target species selection***

From the plant species database, we further classified indicator species for degenerated sites and donor sites based on the Flora of China [21]. *Artemisia frigida*, *Cleistogenes squarrosa*, *Carex duriuscula* and *Potentilla acaulis* were selected as indicator species for degenerated sites. *Leymus chinensis* and xerophytic grasses (*Stipa capillata* and *Koeleria cristata*) were selected as meadow steppe target species. *L*. *chinensis* and mesophytic forbs (*Sanguisorba officinalis*, *Artemisia tanacetifolia*, *Vicia sepium*, *Trifolium lupinaster*, *Lathyrus quinquenervius*, *Galium verum* and *Carex pediformis*) were selected as upland meadow target species [42 – 44].

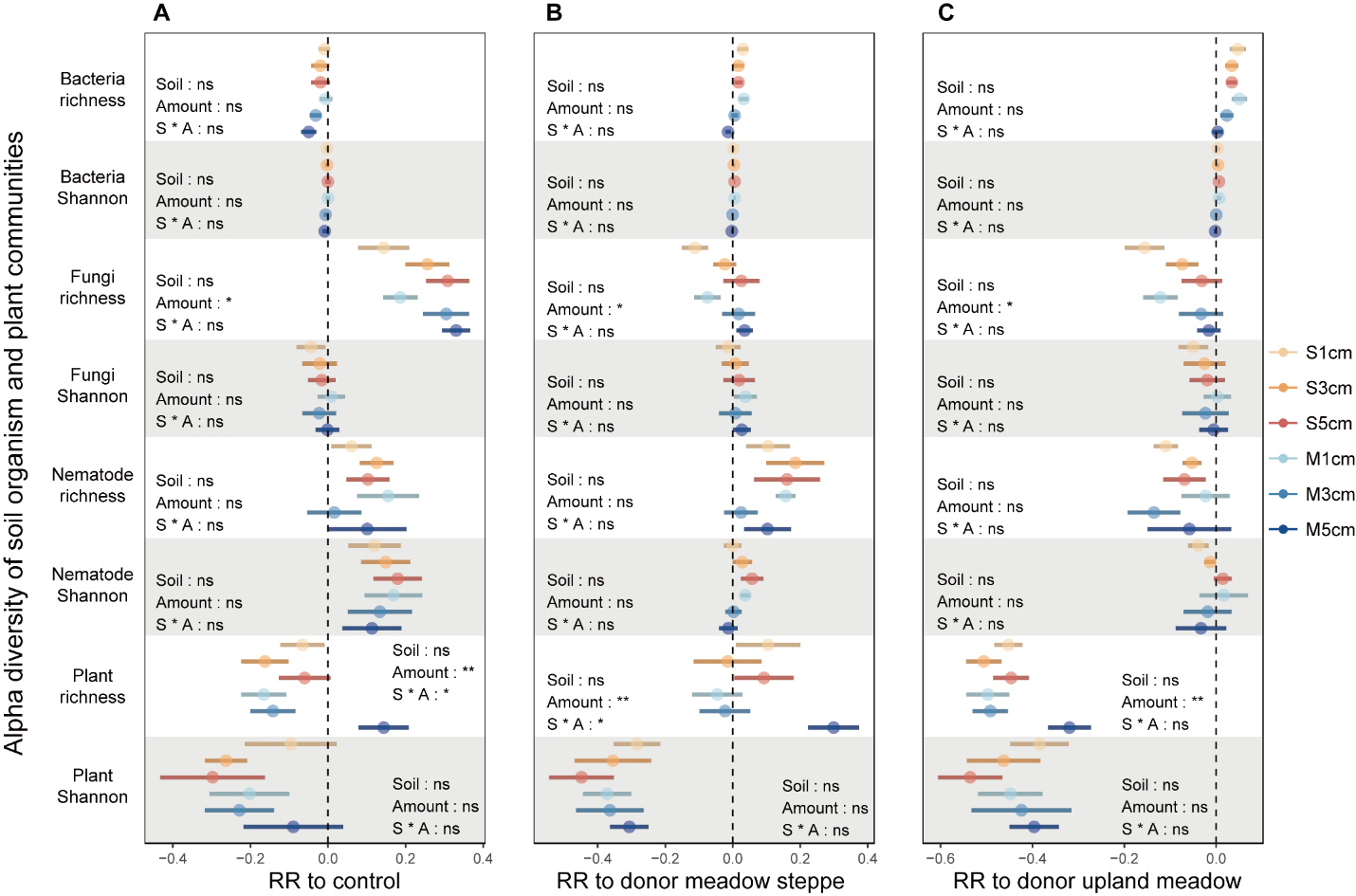

**Figure S1. Response ratio of alpha diversity of soil organism and plant communities to control, donor meadow steppe and donor upland meadow.** Error bars represent ± SE. The category definitions are as follows: S 1cm, meadow steppe soil inoculation with 1 cm depth; S 3cm, meadow steppe soil inoculation with 3 cm depth; S 5cm, meadow steppe soil inoculation with 5 cm depth; M 1cm, upland meadow soil inoculation with 1 cm depth; M 3cm, upland meadow soil inoculation with 3 cm depth; M 5cm, upland meadow soil inoculation with 5 cm depth. * *P* < 0.05; ** *P* < 0.01; *** *P* < 0.001.

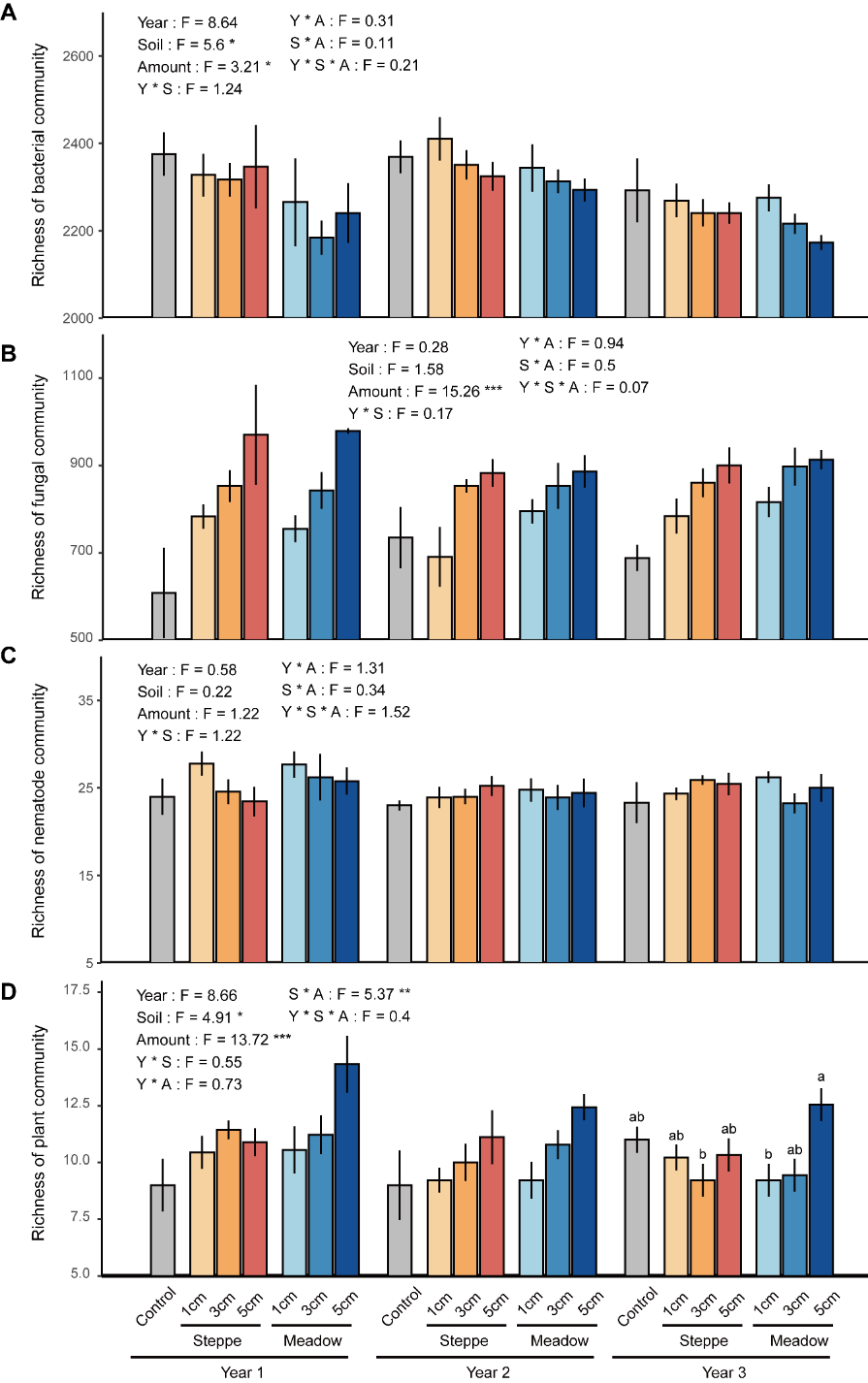

**Figure S2. Soil inoculation effect on the richness of bacterial, fungal, nematode and plant communities.** The numbers at the top represent the result of linear mixed models. * *P* < 0.05; ** *P* < 0.01; *** *P* < 0.001. The category definitions are as follows: Control, the top soil removed treatment (TSR); Steppe, inoculation with meadow steppe soil; Meadow, inoculation with upland meadow soil. Letters in bars represent the difference within one year, different letters indicate significant differences at *P* < 0.05. Error bars represent ± SE.

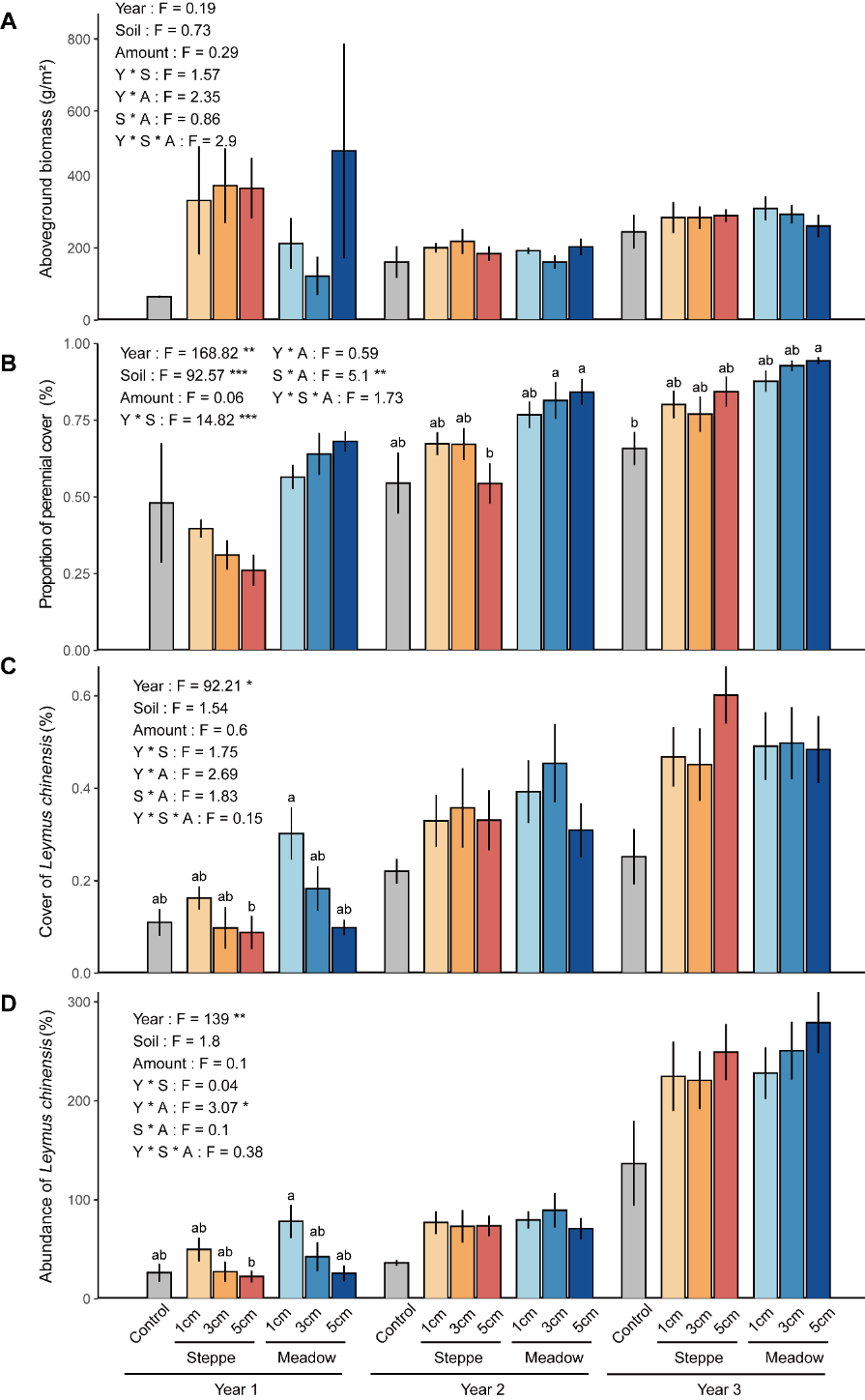

**Figure S3. Soil inoculation effect on plant traits in soil inoculation treatments.** The numbers at the top represent the result of linear mixed models. * *P* < 0.05; ** *P* < 0.01; *** *P* < 0.001. The category definitions are as follows: Control, the top soil removed treatment (TSR); Steppe, inoculation with meadow steppe soil; Meadow, inoculation with upland meadow soil. Letters in bars represent the difference within one year, different letters indicate significant differences at *P* < 0.05. Error bars represent ± SE.

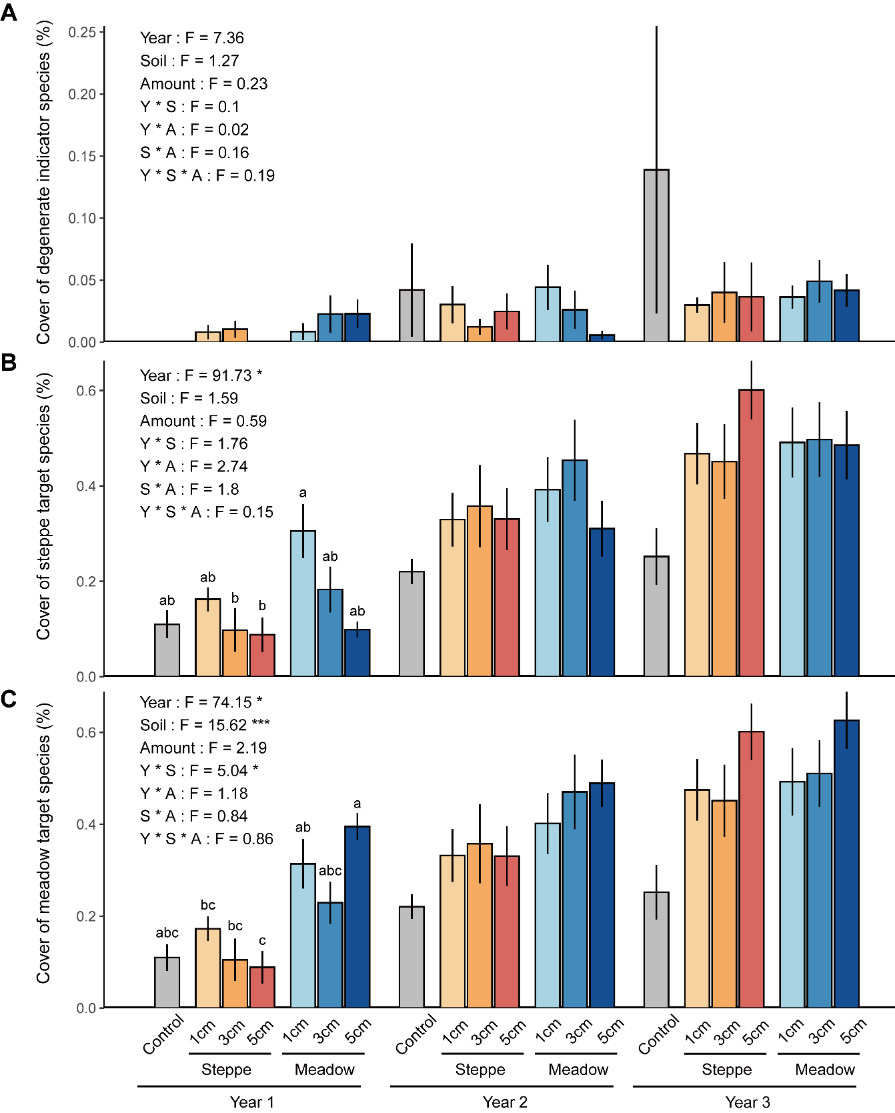

**Figure S4. Soil inoculation effect on the cover of degenerate indicator species, meadow steppe target species and upland meadow target species.** The numbers at the top represent the result of linear mixed models. * *P* < 0.05; ** *P* < 0.01; *** *P* < 0.001. The category definitions are as follows: Control, the top soil removed treatment (TSR); Steppe, inoculation with meadow steppe soil; Meadow, inoculation with upland meadow soil. Letters in bars represent the difference within one year, different letters indicate significant differences at *P* < 0.05. Error bars represent ± SE.

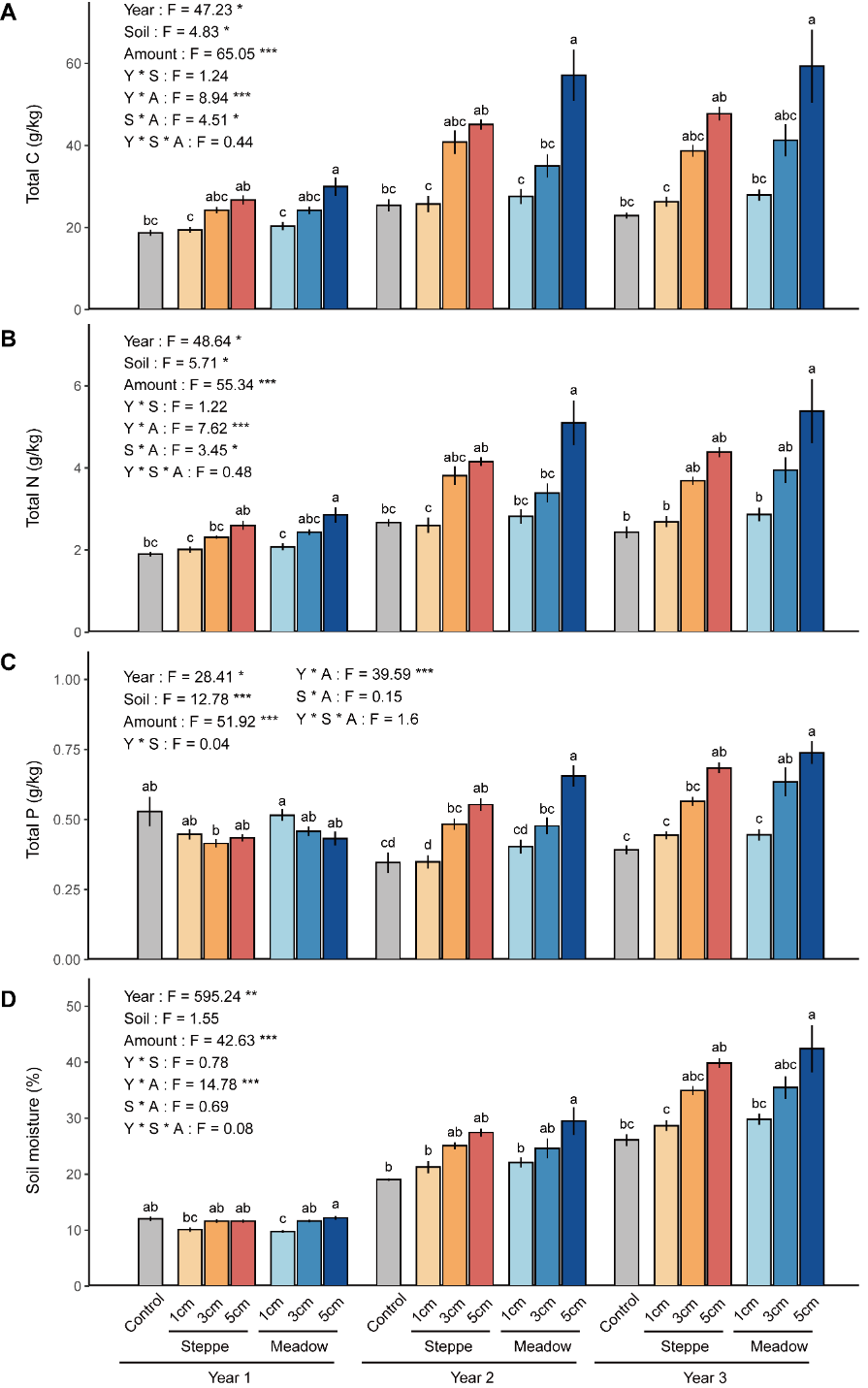

**Figure S5. Soil inoculation effect on soil nutrient and soil moisture.** The numbers at the top represent the result of linear mixed models. * *P* < 0.05; ** *P* < 0.01; *** *P* < 0.001. The category definitions are as follows: Control, the top soil removed treatment (TSR); Steppe, inoculation with meadow steppe soil; Meadow, inoculation with upland meadow soil. Letters in bars represent the difference within one year, different letters indicate significant difference at *P* < 0.05. Error bars represent ± SE.

**Table S1.** Adonis test of the effect soil type, amount and their interactions on soil organism and plant communities in separate years.

|  |  | Year 1 |  | Year 2 |  | Year 3 |  |
| --- | --- | --- | --- | --- | --- | --- | --- |
|  |  | F | *P* | F | *P* | F | *P* |
| Bacteria | Soil | 1.833 | 0.001 | 5.304 | 0.001 | 6.281 | 0.001 |
|  | Amount | 1.504 | 0.002 | 2.108 | 0.001 | 2.214 | 0.001 |
|  | Soil*Amount | 1.257 | 0.058 | 1.326 | 0.053 | 1.539 | 0.010 |
| Fungi | Soil | 1.770 | 0.001 | 4.518 | 0.001 | 4.719 | 0.001 |
|  | Amount | 1.281 | 0.038 | 1.441 | 0.019 | 1.478 | 0.006 |
|  | Soil*Amount | 1.169 | 0.126 | 1.257 | 0.081 | 1.170 | 0.118 |
| Nematodes | Soil | 1.305 | 0.151 | 3.239 | 0.001 | 8.438 | 0.001 |
|  | Amount | 0.981 | 0.496 | 1.593 | 0.028 | 1.611 | 0.049 |
|  | Soil*Amount | 0.670 | 0.920 | 0.800 | 0.779 | 1.637 | 0.051 |
| Plants | Soil | 15.463 | 0.001 | 7.258 | 0.001 | 5.958 | 0.001 |
|  | Amount | 5.187 | 0.001 | 4.198 | 0.001 | 2.534 | 0.001 |
|  | Soil*Amount | 3.119 | 0.001 | 2.071 | 0.008 | 1.305 | 0.157 |

**Table S2.** Topological features of the co-occurrence networks of empirical networks and random networks.

|  | Empirical network | |  | Random network | |
| --- | --- | --- | --- | --- | --- |
|  | Meadow steppe | Upland meadow | | Meadow steppe | Upland meadow |
| Nodes | 212 | 266 | | 212 | 266 |
| Edges | 329 | 496 | | 329 | 496 |
| Complexity | 1.55 | 1.86 | | 1.55 | 1.86 |
| Clustering coefficient | 0.26 | 0.3 | | 0.0029 | 0.011 |
| Centralization closeness | 0.0072 | 0.0044 | | 0.022 | 0.016 |
| Centralization betweenness | 0.22 | 0.11 | | 0.081 | 0.056 |
